# Pericyte Heterogeneity Identified by 3D Ultrastructural Analysis of the Microvessel Wall

**DOI:** 10.1101/2022.08.08.503052

**Authors:** Hanaa Abdelazim, Laura Beth Payne, Kyle Nolan, Karan Paralkar, Vanessa Bradley, Ronak Kanodia, Rosalie Gude, Rachael Ward, Aboozar Monavarfeshani, Michael A. Fox, John C. Chappell

**Author notes:** **Corresponding Author and Lead Contact:** John C. Chappell, Ph.D., Associate Professor, Fralin Biomedical Research Institute at Virginia Tech-Carilion, 2 Riverside Circle, Roanoke, Virginia 24016, E-mail Correspondence. These authors contributed equally to this work (Hanaa Abdelazim, Laura Beth Payne, Kyle Nolan). Previous affiliation(s) (Karan Paralkar, Vanessa Bradley, Rachael Ward, Aboozar Monavarfeshani).

## Abstract

Unequivocal pericyte identification remains a limitation in the field of vascular biology given the lack of unique molecular marker. Compounding this challenge are the recently described heterogeneities in pericyte morphology across microvascular networks. Here, we found further support on the ultrastructural level for classifying pericytes into sub-types – “thin-strand” (TSPs), mesh (MP), and ensheathing (EP) pericytes – based on their architecture in the mouse brain microcirculation. We also observed several instances of an additional cell type in the medial layer between endothelial cells and pericytes, specifically associated with EPs. A conserved characteristic across PC subtypes was extracellular matrix (ECM) encompassing the vascular unit and dispersed among neighboring cells. ECM thicknesses fell within a specific range depending on vessel location, and only thinned where cells were in closer proximity. Pericytes and endothelial cells formed “peg-and-socket” structures at these locations, providing another distinguishing feature across PC subtypes. Unique contact locations seemed to be present between medial and endothelial cells, as well as between vascular cells and the brain parenchyma. The ECM surrounding EPs exhibited another notable configuration in that thin extensions radiated out from the vessel wall into the surrounding parenchyma, suggesting mechanical and/or biochemical roles. Considering these data together, ultrastructural observations may provide an orthogonal perspective on pericyte heterogeneity and the presence of medial cells in cerebrovascular walls as well as assessing ECM coverage as a criterion for PC identification and exploring PC-associated ECM extensions that may have unique relevance in health and disease.

## INTRODUCTION

Pericytes (PCs) are essential cellular components of the capillary wall, maintaining microvascular integrity in a variety of organs and tissues (Payne et al., 2020), most notably within the blood-brain barrier (BBB) (Daneman et al., 2010). Along with established roles in vascular barrier function, PCs have been assigned additional functions including regulating microvessel tone (Hall et al., 2014; Hartmann et al., 2021; Hill et al., 2015), modulating immune cell trafficking (Navarro et al., 2016; Sava et al., 2015), and phagocytic clearance within the capillary microenvironment (Rustenhoven et al., 2017). To better understand these potential roles for PCs within the microcirculation, a wide range of approaches have been applied to characterize their structure on both the cellular and ultrastructural levels.

Recent studies have revealed that PC morphology is likely far more diverse than their colloquial characterization as “bumps-on-a-log.” Depending on their location within blood vessel networks, PCs can be found with extensive cellular processes covering non-overlapping domains of the capillary endothelium, while others appear to form a “mesh-like” configuration around microvessels (Grant et al., 2019). Approaching transitions to larger diameter vessels such as arterioles and venules, PCs seem to ensheath vessel segments and cover almost the entirety of the abluminal surface with their cellular extensions (Ivanova et al., 2021). Thus, PC structural heterogeneity reflects a similar diversity and specification found with endothelial cells (ECs) (Potente and Makinen, 2017), which can differ in arterio-venous identity, architecture within specialized capillaries (e.g. continuous, fenestrated, or discontinuous), and harbor tissue-specific properties.

Alongside this increased appreciation of PC morphological diversity, several key features seem to be conserved across PC sub-types. For instance, most, if not all, PCs appear to be encased in a specialized extracellular matrix (ECM) known as the vascular basement membrane (vBM), also referred to as the basal lamina if considering the neurovascular unit (Payne et al., 2019; Sava et al., 2015; Stratman et al., 2009). The composition of the vBM may vary depending on the tissue or organ, but PCs and ECs synthesize and deposit a number of ECM components such as various collagens, laminins, and fibronectin (Stratman et al., 2009). The extent to which the vBM thickness may vary across microvessel regions within and between various organs remains to be determined in healthy and diseased states (Lopez-Luppo et al., 2017; Thomsen et al., 2017; Tien et al., 2014; Wimmer et al., 2019). Nevertheless, it is widely accepted that the vBM between PCs and ECs thins and is likely absent at locations of direct PC-EC contact known as “peg-and-socket” junctions (Caruso et al., 2009; Diaz-Flores et al., 1991; Ornelas et al., 2021). These junctions are proposed to be domains for PC-EC coupling via gap junctions (Diaz-Flores et al., 1991; Payne et al., 2022) as well as adhesion protein localization (e.g. N-Cadherin) (Diaz-Flores et al., 2009; Fang et al., 2013; Gerhardt et al., 2000; Tillet et al., 2005). A recent study by Ornelas et al. incorporated focused ion beam-scanning electron microscopy (FIB-SEM) approaches to identify EC protrusions (i.e. “pegs”) extending into adjacent PCs, which were less abundant than the PC extensions towards and into neighboring ECs (Ornelas et al., 2021). Thus, new imaging modalities and experimental techniques will continue to reveal unique aspects of the vBM between microvascular cells, and a broader array of cellular interactions within the capillary wall than previously identified.

Pericytes primarily interact with the endothelium and with adjacent PCs, while their direct interaction with other cell types remains poorly understood. Though PCs may share a common lineage with vascular smooth muscle cells (vSMCs) as seen in the development of the coronary arteries (Volz et al., 2015), it is unclear if PCs and vSMCs remain in direct contact in mature vasculature and within all tissue beds. In the brain, for instance, PC coupling to vSMCs may represent a mechanism by which downstream vasoconstriction/dilation signals can propagate upstream to arteriolar smooth muscle and induce vessel diameter changes (Gonzales et al., 2020; Hall et al., 2014). Furthermore, astrocytes (ASCs) within the brain have also been proposed to directly couple to vascular cells (Petzold and Murthy, 2011). It remains to be determined however if PCs and ASCs form cell-cell contacts/junctions, or if they are separated by the vBM at all locations and depend solely on paracrine signaling for their communication. Cells involved in immune surveillance have also been proposed to reside adjacent to blood vessels including microglia (Bisht et al., 2021) and perivascular macrophages (Faraco et al., 2017; Faraco et al., 2016), but the extent to which these cells may be located within the vBM, if at all, is still being established.

In the current study, we sought to address some of these open questions regarding PC structural heterogeneity and their relationship with the vBM and surrounding cell types. Specifically, we analyzed and segmented several datasets of mouse brain tissue prepared for and imaged by serial block face-scanning electron microscopy (SBF-SEM). This analysis revealed multiple instances of PCs wrapped around microvessels with a single cell in the medial layer, presumably a vSMC, which contacted the underlying endothelium via unique cellular structures. These ensheathing PCs, corroborated by high-power confocal imaging, were distinct in their ultrastructure from other microvascular PCs. These other PC subtypes appeared to be configured with either “mesh-like” projections or single cellular extensions (i.e. “thin-strand”) along brain capillaries. We found the vBM within all regions to be remarkably consistent in thickness except where direct PC-EC contact points occurred. We also observed ASC end-feet along the vessel wall, with few obvious locations where direct ASC-PC coupling could be identified. Interestingly, we found distinct bands of ECM extending from ensheathing PCs into surrounding ASCs, which were absent along thin-strand and mesh-type PCs.

## MATERIALS AND METHODS

### Serial Block Face-Scanning Electron Microscopy Imaging of Mouse Brain Tissue

#### Tissue Preparation and SBF-SEM Imaging Parameters

All animal experiments were conducted with review and approval from Virginia Tech Institutional Animal Care and Use Committee (IACUC), which reviewed and approved all protocols. The Virginia Tech NIH/PHS Animal Welfare Assurance Number is A-32081-01 (expires July 31, 2025). Methodology for generating SBF-SEM datasets can also be found in Hammer et al. 2015 (Hammer et al., 2015). Briefly, wild-type C57BL/6 mice were obtained from Charles River, and Lrrtm1^−/−^ mice were acquired from MMRRC (stock #031619-UCD). Mice were trans-cardially perfused sequentially with PBS and 4% paraformaldehyde/2% gluteraldehyde in 0.1M cacodylate buffer. Brains were immediately removed and sectioned by vibratome (300 μm coronal sections), and the dorso-lateral geniculate nucleus (dLGN) were dissected. Tissues were further processed, embedded, sectioned, and imaged by Renovo Neural Inc. Images were acquired at a resolution of 5 nm/pixel, and image sets included >200 serial sections (with each section representing 75 nm in the z axis). SBFSEM data sets were 40 μm × 40 μm × 12–20 μm.

#### Image Processing and Segmentation, and Vessel Morphology Quantification

A total of 24 tissue sample image sets were generated with a sum of 12,675 images. To optimize image analysis, we filtered images by selecting ones containing vessels of interest, lowering the number of images to 7,726. A sizeable number of remaining images were not analyzable due to poor resolution, contamination of artifacts or poor vessel orientation (i.e. longitudinal vs. cross-sectional). Thus, 3,494 images remained and were deemed suitable for analysis. Each image stack was loaded into FIJI/ImageJ, and vessels of interest were identified based cellular localization around the vessel lumen and morphologies. Cellular segmentation was performed via the TrakEM2 plugin for FIJI. Pericyte circumferential coverage was assessed by first measuring the vBM surrounding the ECs to determine the circumference of the vessel. The inner vBM of each PC was then measured, and the percent of shared vBM between PC and EC was reported. We averaged five random locations of vBM between structures of interest to assess vBM thickness. Ten images, equally spaced throughout the image stack (i.e. along the z-axis), were assessed for circumferential coverage and vBM thickness. Areas containing peg-and-socket junctions, where vBM thickness was considerably smaller, were omitted from this portion of the analysis.

### Confocal Imaging of Mouse Brain Tissue

#### Tissue Preparation

Adult WT mouse brains were extracted from animals following trans-cardial perfusion of PBS and subsequent 4% paraformaldehyde in PBS. Brains were immersed in a 30% sucrose solution overnight to provide cryo-protection for subsequent slicing by cryostat at a thickness of 10-20 μm.

#### Immunofluorescence Labeling

Sections on slides were encircled with a hydrophobic solution and incubated with a blocking solution containing 5% serum and 0.1% Triton-X. Following PBS washes, primary antibodies were sequentially added with washing steps in between, specifically: goat anti-mouse platelet-endothelial cell adhesion molecule-1 (PECAM-1)/CD31 (R&D Systems), mouse anti-mouse α-smooth muscle actin (αSMA) conjugated to Cy3 fluorophore (Sigma), and mouse anti-rat neural-glial antigen 2 (NG2) (Abcam). Appropriate mouse-on-mouse blocking solutions were applied to prevent non-specific binding of secondary antibodies. Secondary antibodies applied included donkey anti-goat IgG conjugated to AlexaFluor647 (Jackson ImmunoResearch) and donkey anti-mouse IgG1 conjugated to AlexaFluor488 (Jackson ImmunoResearch).

#### Confocal Imaging Parameters

Confocal images were taken on a Zeiss LSM880 confocal microscope using a 100x water-immersion objective. Images were taken through the thickness of the slice, and spacing for z-stack images was optimized for 50% overlap in adjacent optical slices.

### Statistical Analysis

GraphPad Prism 8 software was used for statistical analysis. For measurements where statistical comparisons are shown, we applied an ordinary one-way Analysis of Variance (ANOVA) test followed by a Tukey’s multiple comparisons test to detect differences between each group. Statistical significance was set at a P value less than or equal to 0.05. Measurements were taken across a minimum of n=3 for vessel segments associated with each PC subtype (TSP, MP, and EP), with technical replicate measurements taken where possible for each vessel replicate.

## RESULTS

### Three Pericyte Subtypes Displayed Distinct Ultrastructural Morphologies

Several molecular and morphological criteria have been proposed for identifying microvascular PCs (Armulik et al., 2011). Pericytes deep within capillary networks are frequently described as having prominent somas containing their nuclei, with projections emanating along the vessel wall (Ivanova et al., 2021). These vascular mural cells harboring narrow cellular extensions recently garnered the designation of “thin-strand” PCs (TSPs) (Grant et al., 2019). We focused our initial analysis of SBF-SEM images from mouse brain tissue on these TSPs, finding numerous instances of PCs with this configuration (Figure 1A). TSPs exhibited discrete nuclei and cytoplasm largely separated from adjacent cells by a thin layer of ECM. Capitalizing on the volumetric perspective afforded by SBF-SEM datasets, we were able to track the presence of TSP cytoplasmic projections along the abluminal surface of the capillary wall. These extensions narrowed substantially as they progressed along the microvessel, and three-dimensional (3D) rendering of segmented datasets revealed morphology consistent with the “thin-strand” designation (Figure 1A, see Supplemental Movie 1) derived from optical imaging (Grant et al., 2019).

**Figure 1.**
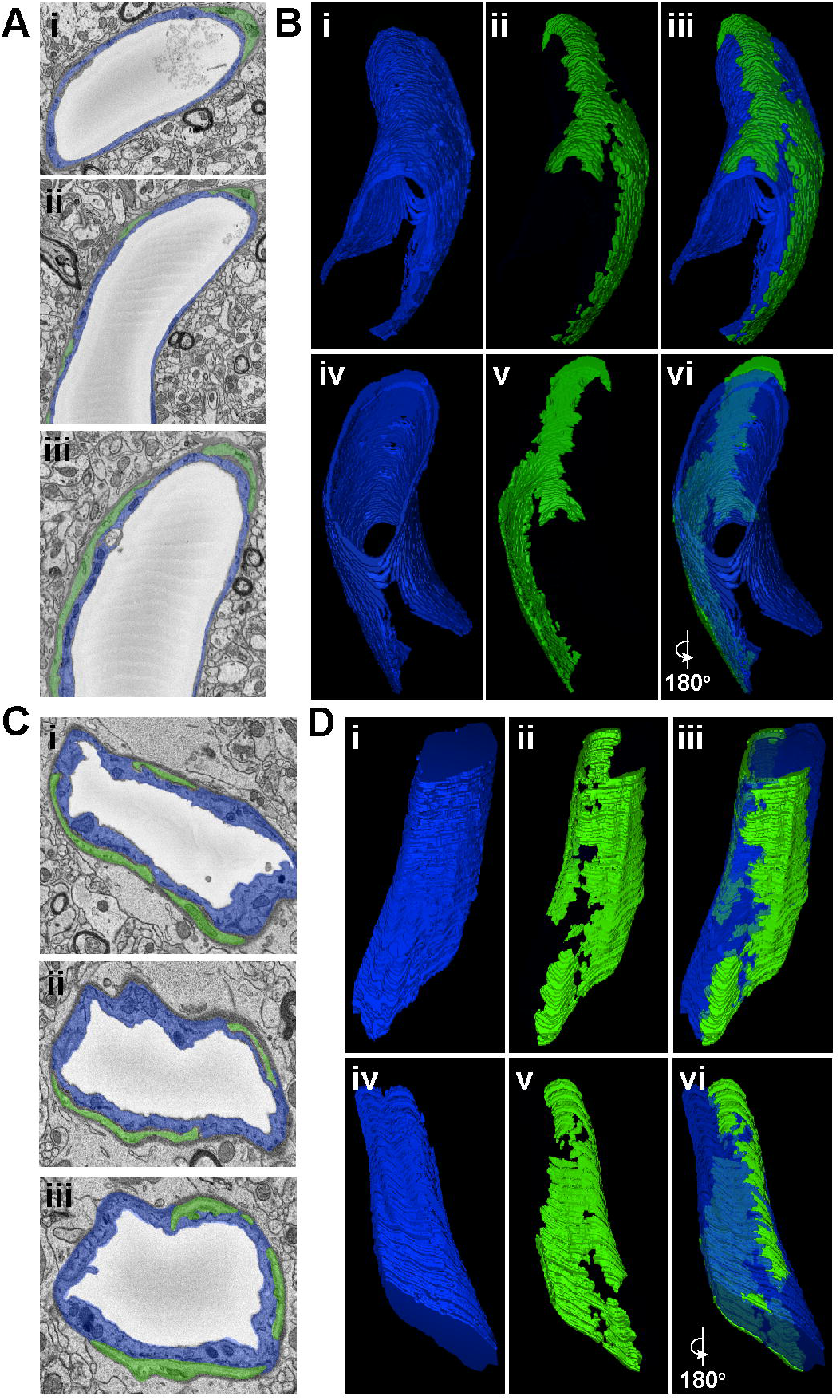
Representative SEM Images and 3D Renderings of Thin-Strand and Mesh Pericytes in Mouse Brain Microvessels. (A) SEM images of an EC (pseudo-colored blue in i, ii, and iii) and thin-strand PC (pseudo-colored green in i, ii, and iii) within the wall of a mouse brain microvessel. (B) 3D renderings of an EC (blue in i, iii, iv, and vi) and a thin-strand PC (green in ii, iii, v, and vi) generated from annotated SBF-SEM datasets. To allow for a more complete view of the vessel, EC renderings have a reduced opacity, and images in iv, v, and vi are a 180-degree rotation of images in i, ii, and iii. *See corresponding Supplemental Movie 1*. (C) SEM images of an EC (pseudo-colored blue in i, ii, and iii) and mesh PC (pseudo-colored green in i, ii, and iii) within the wall of a mouse brain microvessel. (D) 3D renderings of an EC (blue in i, iii, iv, and vi) and a mesh PC (green in ii, iii, v, and vi) generated from annotated SBF-SEM datasets. To allow for a more complete view of the vessel, EC renderings have a reduced opacity, and images in iv, v, and vi are a 180-degree rotation of images in i, ii, and iii. *See corresponding Supplemental Movie 2*.

Previous studies have described another subset of PCs displaying a “mesh-like” configuration of their cellular processes along the microvessel wall. These distinct mesh PCs (MPs) were present in our SBF-SEM datasets as well, facilitating their ultrastructural characterization. Similar to TSPs, MP nuclei and associated cytoplasm occupied abluminal positions beyond the vBM adjacent to ECs but still contained within a layer of ECM separating the vessel unit from surrounding parenchyma (Figure 1B, see Supplemental Movie 2). Volumetric analysis however revealed features contrasting with TSPs, specifically that MP cytoplasm was present along a greater extent of the microvessel surface. Again, the power of 3D ultrastructural rendering underscored this distinction from TSPs (Figure 1B), further supporting the notion that, within the brain tissue analyzed in the current study, PCs could be classified into subsets based on their ultrastructural morphology.

While TSPs and MPs appeared to be the more prevalent PC subtypes along brain capillaries (Figure 2A), we found another perivascular cell exhibiting features consistent with a PC subpopulation previously suggested to surround the basolateral surface with a thin and nearly continuous layer of cytoplasm (Grant et al., 2019). This PC subtype has been described as “ensheathing” PCs (EPs). These EPs occupied a greater percentage of the vessel circumference as compared to MPs and TSPs (Figure 2B, see Supplemental Movie 3), with more cross-sectional area associated with the underlying ECs (Figure 2C). MPs displayed an intermediate phenotype with respect to these measurements, and TSP had the lowest circumferential distribution and cross-sectional area quantified (Figure 2B-C). Interestingly, we found several instances in which a third cell type was present between the EC layer and the outermost PC layer. These medial cells were enriched for mitochondria, and their nuclei were oriented circumferentially, consistent with hallmarks of vSMCs (Park et al., 2014; Thakar et al., 2009) (Figure 2D), though the inability to apply cell-specific labeling approaches limited a more precise classification. Nevertheless, we applied high-resolution confocal imaging to a separate set of immunostained mouse brain sections to aide in the interpretation of the identity of these medial cells (Figure 2E). We found several instances of cell nuclei strongly associated with: (i) an outmost vessel layer positive for neural-glial antigen-2 (NG2), an accepted marker of brain PCs, (ii) a medial cell layer containing signals for α-smooth muscle actin (αSMA), a contractile protein highly enriched in vSMCs, and (iii) an innermost EC layer labeled for platelet-endothelial cell adhesion molecule-1 (PECAM-1/CD31) (Figure 2E). Taken together, these data suggest that three distinct PC subpopulations may be identified within the brain microcirculation by their ultrastructural characteristics, and that EPs cover microvessel regions containing phenotypically distinct mural cells.

**Figure 2.**
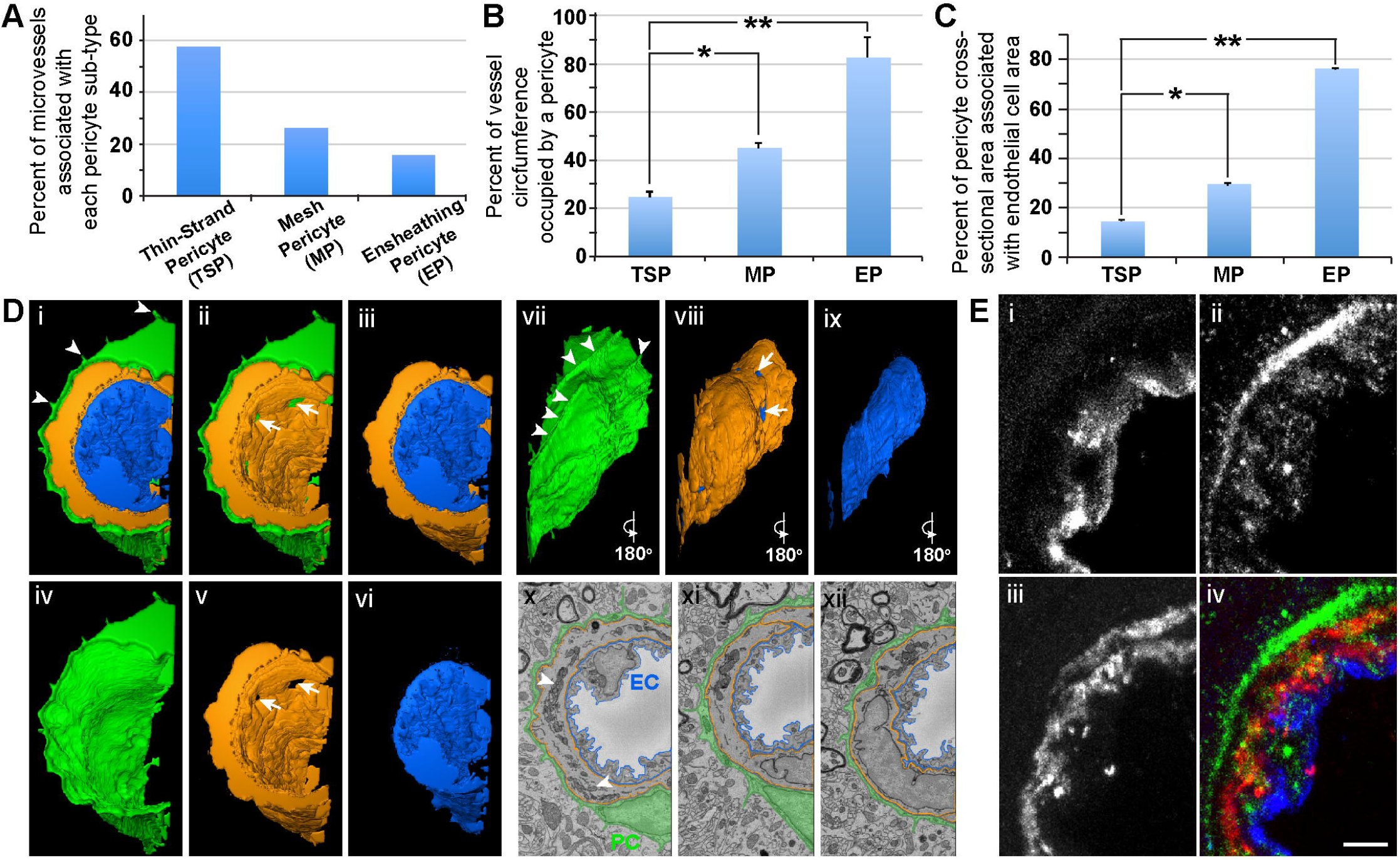
Quantitative Comparison of PC Subtypes and Representative SEM Images, 3D Rendering, and Optical Images of an Ensheathing PC in Mouse Brain Microvasculature. (A) Bar graph displaying the percent of microvessels associated with each PC subtype: thin-strand PC (TSP), mesh PC (MP), and ensheathing PC (EP). (B) Bar graph displaying the average percent of the vessel circumference occupied by PCs for each PC subtype. Error bars denote standard deviations. *P≤0.05 and **P≤0.01 for the comparisons shown. (C) Bar graph displaying the average percent of cross-sectional area of each PC subtype associated with underlying EC cross-sectional area. Error bars denote standard deviations. *P≤0.05 and **P≤0.01 for the comparisons shown. (D) 3D renderings and SEM images of an EC (blue in i, iii, vi, ix, x-xii), an ensheathing PC (green in i, ii, iv, vii, x-xii), and a medial cell (orange in i, ii, iii, v, viii, x-xii) generated from annotated SBF-SEM datasets. Arrowheads in (i and vii) denote unique cellular extensions from EPs associated with ECM extensions. Arrows in (ii, v, and viii) denote gaps in the medial cell allowing a PC-EC interface. To allow for a more complete view of the vessel, individual renderings are shown (iv, v, and vi), and images in vii, viii, and ix are a 180-degree rotation of images in iv, v, and vi, respectively. *See corresponding Supplemental Movie 3*. SEM images of an EC (pseudo-outlined blue in x-xii), ensheathing PC (pseudo-colored green in x-xii), and a medial cell (pseudo-outlined in x-xii) within the wall of a mouse brain microvessel. Arrowheads denote EM signatures consistent with mitochondria in the medial cell. (E) Representative confocal images of mouse brain vessel immunolabeled for Platelet-Endothelial Cell Adhesion Molecule-1 (PECAM-1, i and blue in iv), Neural-Glial Antigen 2 (NG2, ii and green in iv), and α-Smooth Muscle Actin (αSMA, iii and red in iv). Scale bar, 2 μm.

### The Microvascular Basement Membrane is Maintained within a Narrow Range of Thickness Between Vascular Cells and at the Parenchyma Interface

Another essential component of the microvessel wall is the vBM, a thin layer of ECM that separates vascular cells from each other and from the surrounding parenchyma (Payne et al., 2019; Sava et al., 2015; Stratman et al., 2009). The subcellular resolution of our SBF-SEM datasets allowed us to ask if the vBM associated with each PC subtype also displayed distinct features corresponding to their microvascular region. Drawing from our morphological data, we constructed working models for each PC configuration within the blood vessel wall (Figure 3A) and assessed the vBM thickness between each cellular compartment (Figure 3B). Within TSP vessel regions, the vBM thickness at the PC-EC interface was remarkably consistent, with a mean of 150 nm (Figure 3C). The thicknesses between parenchymal cells and each of these vascular cells were also within a narrow range, averaging about 200 nm. The vBM between MPs and ECs was slightly thicker than for TSPs; however, the ECM thickness between the parenchyma and MPs, and relative to their associated ECs, was greater, with a mean of about 250 nm (Figure 3C). EPs displayed the thinnest vBM between themselves and parenchymal cells, less than that of TSPs and significantly thinner than for MPs. In contrast, the medial cells underlying EPs had, on average, the thickest ECM between themselves and the surrounding parenchyma, almost reaching 300 nm in thickness (Figure 3C). These medial cells were also separated from ECs and EPs with ECM akin to the internal elastic lamina (IEL) of arteries, which ranged from 220 nm to 250 nm thick. These data suggest that the vBM encasing the mouse brain microvessels analyzed in the current study is deposited within a well-defined range of thicknesses between each cellular constituent and at the vessel-parenchyma interface, and these ECM densities appear to correspond to microvessel region and PC subtype.

**Figure 3.**
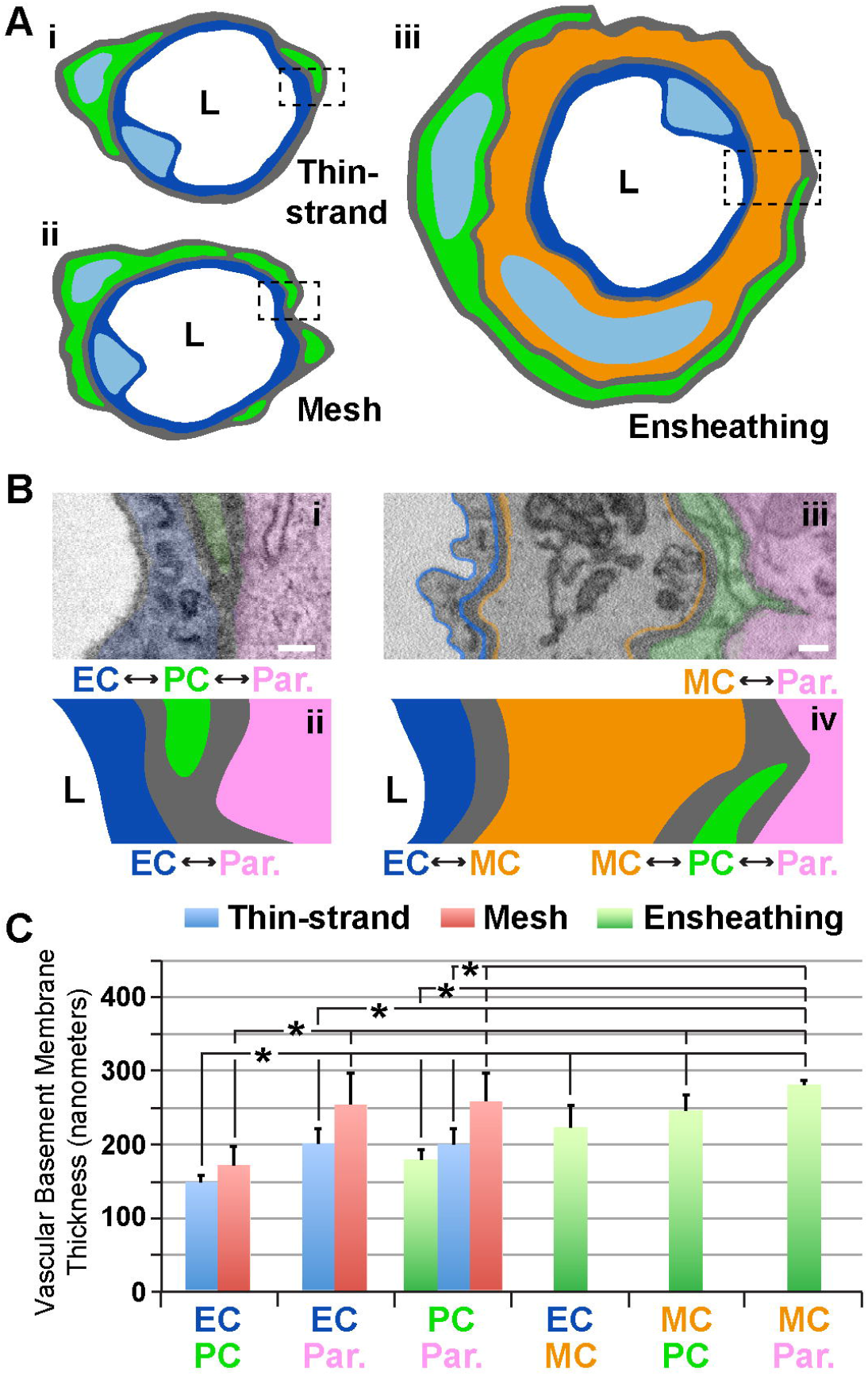
Illustrative Schematics and SEM Images of Vessel Walls Associated with Each PC Subtype and Quantification of ECM Thickness at Each Cellular Interface. (A) Simplified schematics of PCs (green) classified as thin-strand (i), mesh (ii), or ensheathing (iii), with associated endothelium (blue in i-iii) and a medial cell (orange in iii). “L” denotes the vessel lumen. Dashed rectangles indicate the type of vessel cross-section shown in (B). (B) Annotated SEM images illustrating the relative location of the endothelium (blue, pseudo-color in i and drawn in ii, pseudo-outline in iii and drawn in iv), PCs (green, pseudo-colored in i and iii, and drawn in ii and iv), brain parenchyma (pink – Par., pseudo-colored in i and iii, and drawn in ii and iv), and a medial cell (orange – MC, pseudo-outline in iii and drawn in iv). “L” denotes the lumen side. Each interface is denoted with double-sided arrows. Scale bars, 500 nm. (C) Bar graph displaying the vascular basement membrane thickness in nanometers for each of the types of cell-cell interface. Blue bars represent averages for TSPs, red bars represent averages for MPs, and green bars represent averages for EPs. Error bars denote standard deviations. *P≤0.05 for the comparisons shown.

### The Density and Configuration of Presumptive Contact Points between Brain Microvascular Cells Appears to be Region-Specific

As discussed above, we observed that the ECM separating each microvascular cell had a relatively uniform thickness along a given vessel length. At discrete locations between each cell, however, the ECM would thin considerably. ECM thinning in these regions appeared consistent with adjacent cell membranes physically engaging, particularly ECs and PCs, which are known to form “peg-and-socket” domains (Caruso et al., 2009; Diaz-Flores et al., 1991; Ornelas et al., 2021). These unique structures have been described as enriched for adherens junctions e.g. N-Cadherin (Gerhardt et al., 2000) and for gap junction localization to facilitate direct cellular communication (Diaz-Flores et al., 1991; Payne et al., 2022). Until recently, pericyte-derived “pegs” extending towards EC “sockets” were thought to be the predominant configuration (Zhao et al., 2015); however, EC pegs extending into PC sockets have also been described (Ornelas et al., 2021), and were observed in our datasets as well. Analyzing our SBF-SEM images for both types of “peg-and-socket” structures, we sought to address the hypothesis that their density per vessel length likely varies among PC subpopulations. For TSPs, we found an average of approximately one (1) “peg-and-socket” junction between PCs and ECs per micron of vessel length (Figure 4A, see Supplemental Movie 4). Mesh PCs had a higher density of these domains with almost three (3) “peg-and-socket” junctions per micron of vessel length (Figure 4B). Surprisingly, we found minimal to no “peg-and-socket” junctions between EPs and adjacent medial cells, or between EPs and underlying ECs (Figure 4C & D). These data suggest that the density of “peg-and-socket” junctions along a microvessel length likely corresponds to PC subtype and location within the microvasculature.

**Figure 4.**
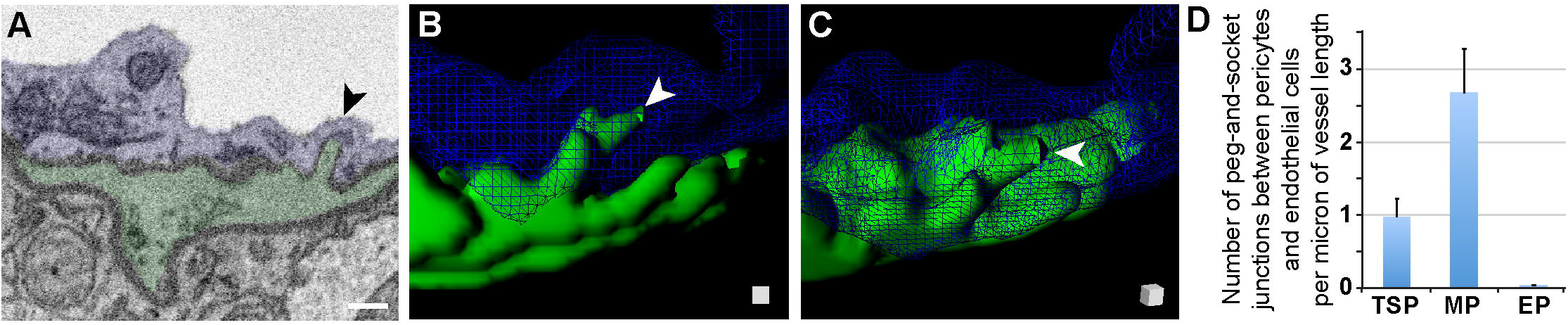
Peg-and-Socket SEM Images, 3D Rendering, and Quantification from Mouse Brain Microvessels. (A) Representative SEM image of a peg-and-socket structure formed by a microvessel EC (blue, pseudo-colored) and adjacent PC (green, pseudo-colored). Arrow denotes the peg-and-socket location. Scale bar, 500 nm. (B) 3D reconstruction of a vessel cross-section with the EC (blue, wire-frame rendering) and PC-derived “peg” (green), noted by a white arrowhead. *See Supplemental Movie 4*. (C) 3D rendering of “peg” shown in (B) with a slight rotation in angle, which is represented by the surface-shaded cube in the lower right corner. “Peg” denoted by white arrowhead. (D) Bar graph displaying the number of peg-and-socket junctions between PCs and ECs per micron of vessel length for each PC subtype. Error bars denote standard deviations.

While EPs in our datasets appeared to lack “peg-and-socket” junctions with neighboring cells, their corresponding medial and endothelial cells did form distinct junctions with one another. As discussed above, “peg-and-socket” junctions between PCs and ECs typically appeared as a thin but notable cellular extension from one cell (i.e. “peg”) occupying an invagination in the neighboring cell (i.e. “socket”) (Figure 4B & C). In contrast, medial cell-EC junctions involved a more complex arrangement of cellular extensions such that medial cell processes were extended into and enveloped by associated EC projections (Figure 5A, see Supplemental Movies 5 & 6), aligning with previous reports of endothelial projections contacting arterial smooth muscle (Maarouf et al., 2017). As with more traditional “peg-and-socket” junctions, there did not appear to be a substantial concentration of ECM within these unique domains. Medial cell-EC junctions were less abundant relative to PC-EC “peg-and-socket” junctions (Figure 5C); however, they occupied a greater distance along vessels compared to “peg-and-socket” junctions, which tended to be more compact. Interestingly, these more distinct junctional configurations were not found in between any other cells analyzed in the current study. Overall, our observations support the notion that, within the walls of brain microvessels analyzed in the current study, the density and structure of presumptive contact points where the ECM thins considerably between microvascular cells may be unique to specific regions.

**Figure 5.**
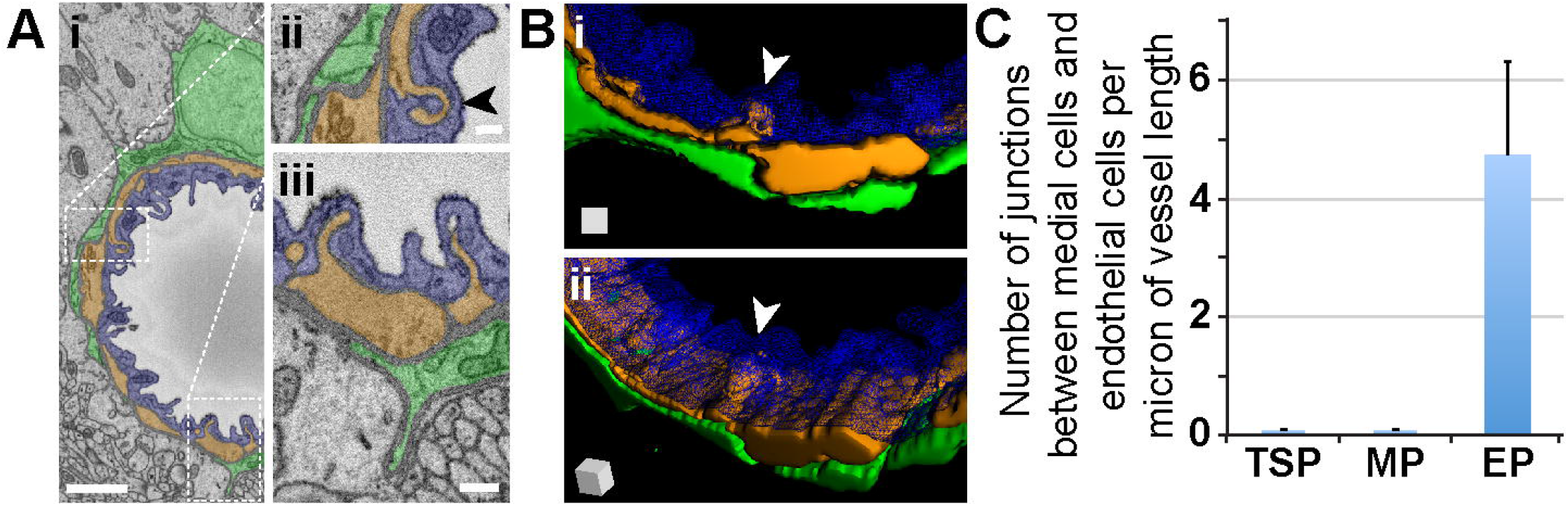
Medial Cell-Endothelium Interface in SEM Images and 3D Renderings, and Quantified from Mouse Brain Microvessels. (A) Representative SEM images of a medial-endothelial cell structures formed within the mouse brain microvasculature. An EC (blue, pseudo-colored i-iii), adjacent PC (green, pseudo-colored in i-iii), and medial cell (orange, pseudo-colored in i-iii) compose the vessel wall. White dotted rectangles denote wall regions shown in higher magnification images in (ii) and (iii). Arrowhead in (ii) denotes a unique medial cell-EC interface further represented in (B). Scale bars in (i) is 1 μm, and in (ii) and (iii) is 200 nm. (B) 3D reconstruction of a vessel cross-section with the EC (blue, wire-frame rendering in i-ii), medial cell-derived structure (orange in i-ii) noted by a white arrowhead, and an associated EP (green in i-ii). *See Supplemental Movies 5 & 6*. The 3D rendering of the medial-endothelial cell structure shown in (ii) is at a slight rotation in angle, which is represented by the surface-shaded cube in the lower left corner. (C) Bar graph displaying the number of medial cell-EC junctions per micron of vessel length for each PC subtype. Error bars denote standard deviations.

### Direct Cell Contact between Microvascular and Parenchymal Cells Appears Minimal, though Ensheathing PCs Appear to Establish Cellular and ECM-Enriched Extensions into the Surrounding Brain Parenchyma

Junctions between cells of the blood vessel wall have been described across various organs (Johnstone et al., 2009) including in neurological tissues. Brain parenchyma cells have also been suggested to directly couple to vascular cells (Petzold and Murthy, 2011). In addition to analyzing the “peg-and-socket” and medial cell-EC junctions described above, we assessed vessels of interest for potential locations where surrounding parenchymal cells such as astrocytes appeared to engage the microvasculature via ECM thinning and direct cell contact (Figure 6A). We did not find any instances of ECM thinning to the extent observed for the other type of junctions. However, reduced ECM intensities at certain locations suggested that unique interface domains may exist between brain parenchymal cells and the associated vasculature (Figure 6A). We quantified the density of these domains per unit length of vessel and found that they were far less abundant than “peg-and-socket” junctions. Microvessel segments associated with TSPs had a density of these unique interface domains around 0.06 domains per micron of vessel length, meaning one of these domains was observed, on average, every 17 microns of vessel length (Figure 6B), compared to TSP “peg-and-socket” junctions detected every micron of vessel length. Capillaries with adjacent mesh PCs displayed an even lower density of these presumptive vessel-parenchyma interface domains, with about 0.02 domains per micron of vessel length, equivalent to one domain every 50 microns of vessel length (Figure 6B). These domains were not detected for vessels encircled by EPs. Collectively, these observations suggested that, within the datasets analyzed herein, areas of direct contact between brain parenchymal cells and the microvasculature were relatively infrequent.

**Figure 6.**
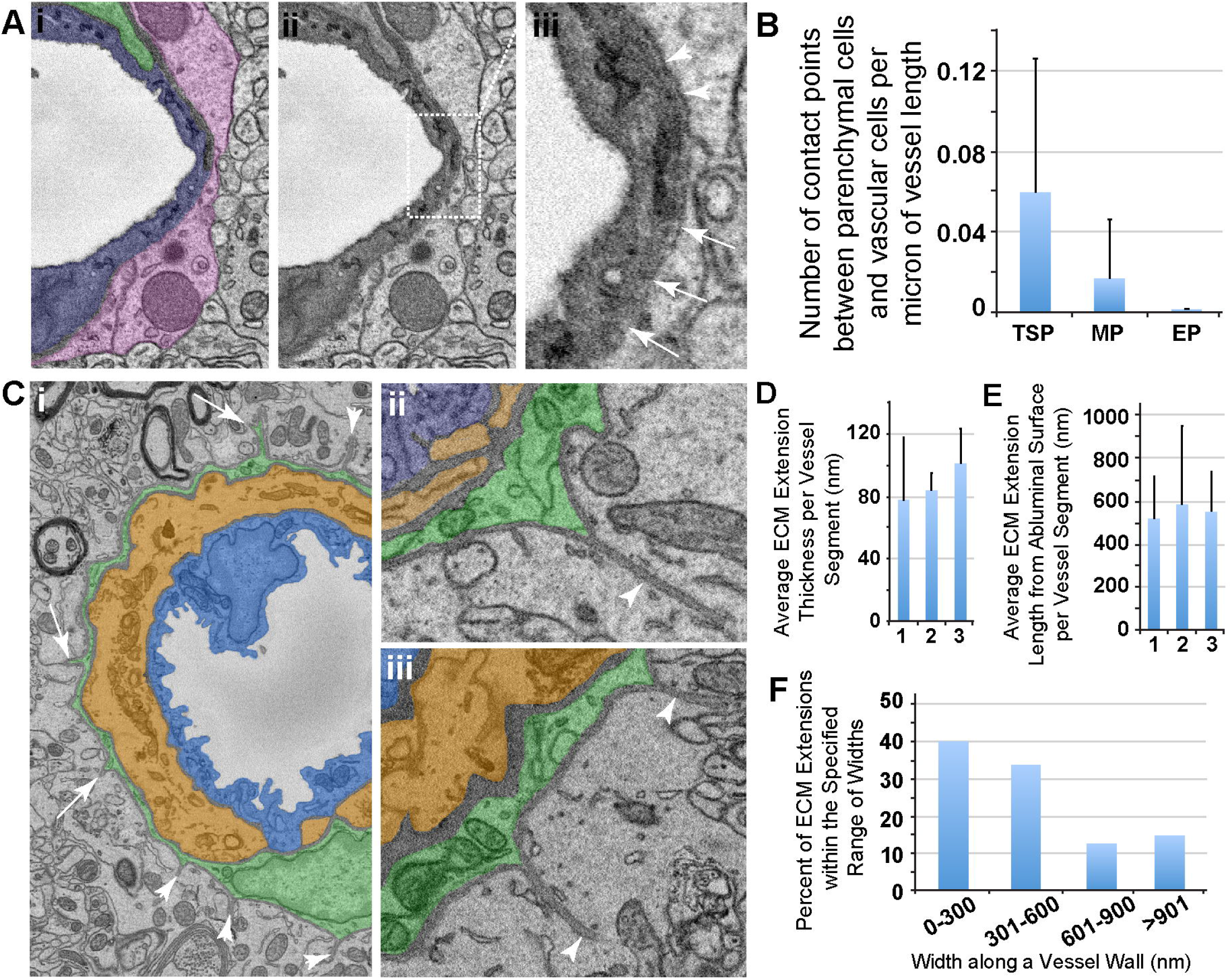
SEM Images and Quantification of Presumptive Parenchyma-Vessel Contact Points and ECM Extensions Emanating from Ensheathing PCs. (A) Representative SEM image of a thin-strand PC (green pseudo-color in i), associated EC (blue pseudo-color in i), and a surrounding parenchymal cell (pink pseudo-color in i). The same image in (i) is shown in (ii) to allow an unobstructed view of the vessel wall SEM. Dotted rectangle denotes a region shown at higher resolution in (iii). The region of interest shown in (iii) represents an area where a denser electron signal was observed along the outer vessel edge (arrowheads), but this signal became thinner along the circumference of the vessel edge (arrows), suggesting the potential for contact between brain parenchyma and the vessel. (B) Bar graph displaying the average number of presumed contact points between brain parenchymal cells and vascular cells per micron of vessel length. Error bars denote standard deviations. (C) Representative SEM images of an ensheathing PC (green pseudo-color in i-iii) with an underlying medial cell (orange pseudo-color in i-iii) and endothelial cell (blue pseudo-color in i-iii). Arrowheads in (i) denote ECM extensions into the surrounding brain parenchyma, and arrows note PC processes associated with ECM extensions found in consecutive images. Images shown in (ii) and (iii) are additional examples of ECM extensions of various lengths, also with associated PC cytoplasm at their base. (D) Bar graph showing the average ECM extension cross-sectional thickness per EP-associated vessel segment (n=3) in nanometers. Error bars denote standard deviations. (E) Bar graph showing the average ECM extension radial length per EP-associated vessel segment (n=3) in nanometers. Error bars denote standard deviations. (F) Percent of ECM extensions within the specified range of widths: 0-300 nm, 301-600 nm, 601-900 nm, and greater than 901 nm. “Widths” represent the distance along the primary vessel axis where an ECM extension was detected.

As we conducted the analysis of potential brain parenchyma-vessel contact sites, we found unique structures associated only with ensheathing PCs. Specifically, we observed ECM configured as extensions projecting away from PCs and into the surrounding brain parenchyma (Figure 6C-E). These extensions appeared contiguous with the vBM, and their dimensions varied substantially along the length of their associated microvessels. For the vessels analyzed, the average ECM extension cross-sectional thickness ranged from around 40 nm to 120 nm (Figure 6G). Their average lengths from the abluminal surface into the brain parenchyma were approximately 300 nm to over 900 nm (Figure 6H). While most of these extensions had a width along the vessel axis within 600 nm, some ECM extensions were beyond this range with a subset wider than 900 nm (Figure 6I). These dimensions suggested an almost “blade-like” configuration of this ECM that was reminiscent of anchoring ECM filaments found on lymphatic vessels (Figure 2, see Supplemental Movie 3). A portion of these ECM structures contained cellular projections from underlying PCs, suggesting that capillary PCs may be contributing to their formation and involved in their function. Nevertheless, these unique ECM extensions associated with EPs may be important for a set of PC functions, and evoke questions about how they are formed and what roles they may play in both normal and pathological microvasculature.

## DISCUSSION

Confident identification of pericytes remains an obstacle in field, as a single molecular marker for these unique perivascular cells remains elusive (Mayr et al., 2021). Adding to this challenge is the recent appreciation that PC populations may be heterogeneous, displaying a range of morphologies within capillary networks (Grant et al., 2019). We found additional support on the ultrastructural level for the classification of these PC subtypes – “thin-strand” (TSP), mesh (MP), and ensheathing (EP) – based on distinct morphological characteristics. Interestingly, we also found several examples of another cell type, likely a vascular smooth muscle cell, in a medial layer between ECs and PCs harboring characteristics of the ensheathing type. A conserved feature across the different PC subtypes was the presence of ECM surrounding the vascular unit and distributed in between neighboring cells (Thomsen et al., 2017). The thickness of this vascular basement membrane was remarkably consistent depending on its location, but never strayed beyond a range of 150-300nm unless thinned to facilitate closer proximity of neighboring cells (suggesting direct contact). The density of PC-EC contact points (“peg-and-socket” structures) was another distinguishing feature across the different PC subtypes (Caruso et al., 2009; Diaz-Flores et al., 1991; Ornelas et al., 2021), as were the apparent contact locations between vascular cells and brain parenchymal cells. In addition to this thinning, the ECM surrounding EPs displayed another unique configuration in the form of extensions that emitted out radially into the surrounding parenchyma. Knowledge of the origin and function of these structures is still emerging, but they appearance suggests the potential for being mechanical elements and/or perhaps signaling nodes via embedded molecular cues. Overall, this unique ultrastructural perspective provides new insights into PC heterogeneity and the presence of medial cells within the microvessel wall, the consideration of ECM coverage as another PC identification criteria, and unique ECM configurations (i.e. radial extensions) that may reveal additional aspects of PC heterogeneity.

Observing the morphology of brain microvasculature by SBF-SEM (Allsopp and Gamble, 1979; Egginton et al., 1996; Ornelas et al., 2021) offers an orthogonal view of PCs and their features that are difficult, to impossible, to observe by other methods such as light microscopy. Ultrastructural approaches have inherent limitations, however, that must be considered when interpreting these data (Bushby et al., 2012). For instance, while we found that TSPs were the most abundant sub-type of PC detected in the available datasets, our analysis was certainly limited in scope given the small tissue volume submitted for SBF-SEM. Nevertheless, of the TSPs observed, they occupied only a small fraction of the capillary circumference, which did not align well with morphological signatures of being inwardly contractile (Hill et al., 2015). Longitudinal contractility may be more reasonable for TSPs, while EPs may be configured more for contraction capable of reducing lumen diameters. Contrary to that interpretation however was the notably thinner cell cytoplasm of EPs along the basolateral vessel surface, as well as the presence of a circumferentially wrapped medial cell containing numerous mitochondria that could potentially generate a contractile force (Park et al., 2014). Furthermore, the EP cytoplasm thickness ranged from 150-300 nm in many locations, and this range is at or below the typical lateral and axial resolution of optical imaging (around for 180-500 nm, at best, for standard confocal microscopes) (Fouquet et al., 2015). This observation evoked a question about these configurations and small dimensions potentially giving the impression that an intracellular marker or cell-surface label within or on a PC might actually be associated with the underlying medial cell, or vice versa. This potential overlap between PCs and these underlying medial cells may be an underappreciated concern and a contributing factor in the at-times confusing results from optical imaging modalities.

As discussed above, we consistently found a medial cell between ensheathing PCs and the underlying endothelium. These cells appeared to be enriched for mitochondria and were circumferentially wrapped (Park et al., 2014; Thakar et al., 2009), though not always continuous, as we found some locations within all microvessels analyzed containing gaps with ECM between medial cell extensions. We could not definitively conclude the specific location of these medial cells with respect to upstream arterioles or downstream venules. But these cells exhibited hallmarks consistent with vascular SMCs identified by immunostaining and confocal microscopy. While vascular SMCs seem to be the most likely cell type present, we cannot fully exclude other possible cell types occupying these locations. Vessel-associated macrophages (VAMs) have been described along brain vasculature, though those appear to reside along larger diameter vessels with little to no association with the microcirculation (Faraco et al., 2017; Faraco et al., 2016). These VAMs have also not been described as circumferentially wrapped but rather extended along the primary vessel axis. This would suggest that the medial cells identified in the current study are unlikely to represent VAMs, but again they cannot be excluded a possible cellular constituent of the wall. Other plausible cell types include microglia (Bisht et al., 2021) and perhaps astrocytes, but these cells typically associate with the cerebrovasculature via cellular extensions or processes (e.g. astrocytic end-feet) without outright incorporation in the vessel wall and within the vBM/basal lamina. In fact, reports of other cell types in perivascular locations have been inconclusive regarding the presence of an outer layer of ECM surrounding these cells, highlighting this as another potentially important criteria for more confident identification of PCs in situ along the microvessel wall.

Throughout the course of this study, we found ECM surrounding most, if not all, microvessels observed. The presence of this ECM in the vBM may be an important consideration for studies attempting to classify and characterize normal and pathological features of PC identity and investment within the vessel wall (Thomsen et al., 2017). We found ECM thicknesses around the different PC subtypes to be maintained in a remarkably narrow range, giving a greater appreciation for the impact that vBM/ECM changes may have in certain disease conditions. A thickening or thinning of the vBM likely has profound consequences for solute and gas exchange across the vessel wall, as well as for how vascular cells interact with their local microenvironment, mechanically and biochemically (Chronopoulos et al., 2011; Lopez-Luppo et al., 2017; Tsilibary, 2003). Locations where the ECM was notably thinner within the vessel wall were associated with cell-cell contact points. Most were affiliated with peg-and-socket junctions (Caruso et al., 2009; Diaz-Flores et al., 1991; Ornelas et al., 2021), which varied in density depending on PC sub-type. Moreover, given the 3D rendering possible with SBF-SEM, the “pegs” that extended from PCs or ECs actually appeared more like a micro-villi, displaying a larger surface area that may be well suited for junction formation and direct cell coupling for exchange or transfer.

Although they did not form morphologically distinct structures, brain parenchymal cells did appear to directly interface and engage with microvascular cells observed in the current study. Compared to peg-and-socket junctions, these locations were relatively rare but ECM thinning seemed to be present at discrete locations where cell-cell contacts may be present. Direct signaling could be feasible in these locations, and could be potentially mediated by secreted or soluble signals exchanged by brain parenchymal cells (e.g. astrocytes, microglia, etc.) and the brain microvasculature (Petzold and Murthy, 2011). Another interesting feature found in this study was the presence of thin ECM extensions into the brain parenchyma, though these were only noted on ensheathing PCs. Given that vessel regions associated with EPs also contained medial cells that may be contractile, it is intriguing to speculate that these ECM extensions may be involved in mechanical anchoring, akin to lymphatic filaments (Leak and Burke, 1968), or perhaps force transmission or sensing. The ECM is also known to harbor biochemical cues such as growth factors that can become tethered to the matrix (Abramsson et al., 2007; Lee et al., 2005), so these extensions may also be signaling nodes. Nevertheless, their function is largely unknown but prompts further studies to determine their origin and physiological roles, if they are remodeled or damaged during disease, and how that might affect capillary perfusion and exchange with surrounding tissues.

In summary, this study has offered additional insights from ultrastructural analysis of mouse brain microvasculature to aide in a better understanding of PC architecture and potential function. PC heterogeneity has been described previously (Grant et al., 2019), and here we found more evidence for distinct features of PC sub-populations. We also observed the presence of a medial cell within the microvessel wall, which may suggest the need to reconsider previous interpretations of PC contractility and the cells that are directly responsible for observed changes in vessel diameter. We provided characterization of the ECM surrounding brain capillaries, noting its (i) consistent dimensions along vessel walls, thinning at discrete locations to enable cell-cell contacts, and (ii) unique configuration as slender extensions into the parenchyma, perhaps for mechanical anchoring. As ultrastructural methods increase in availability, we envision these approaches to be incredibly useful orthogonal techniques in characterizing PCs and their contribution to the development, maturity, and dysfunction of the microvasculature.

## Abbreviations

EC: endothelial cell
PC: pericyte
ECM: extracellular matrix
vBM: vascular basement membrane
TSP: Thin-Strand Pericyte
MP: Mesh Pericyte
EP: Ensheathing Pericyte
SMC: smooth muscle cell
ASC: astrocyte
PECAM-1: Platelet-Endothelial Cell Adhesion Molecule-1
SBF-SEM: serial block face-scanning electron microscopy

## Acknowledgements

We would like to thank all of the members of the Chappell and Fox labs for their support and valued assistance on this project, both materially and intellectually.

## REFERENCES

Abramsson, A., et al., 2007. Defective N-sulfation of heparan sulfate proteoglycans limits PDGF-BB binding and pericyte recruitment in vascular development. Genes Dev. 21, 316–31.

Allsopp, G., Gamble, H. J., 1979. An electron microscopic study of the pericytes of the developing capillaries in human fetal brain and muscle. J Anat. 128, 155–68.

Armulik, A., et al., 2011. Pericytes: developmental, physiological, and pathological perspectives, problems, and promises. Dev Cell. 21, 193–215.

Bisht, K., et al., 2021. Capillary-associated microglia regulate vascular structure and function through PANX1-P2RY12 coupling in mice. Nat Commun. 12, 5289.

Bushby, A. J., et al., 2012. Correlative light and volume electron microscopy: using focused ion beam scanning electron microscopy to image transient events in model organisms. Methods Cell Biol. 111, 357–82.

Caruso, R. A., et al., 2009. Ultrastructural descriptions of pericyte/endothelium peg-socket interdigitations in the microvasculature of human gastric carcinomas. Anticancer Res. 29, 449–53.

Chronopoulos, A., et al., 2011. High glucose-induced altered basement membrane composition and structure increases trans-endothelial permeability: implications for diabetic retinopathy. Curr Eye Res. 36, 747–53.

Daneman, R., et al., 2010. Pericytes are required for blood-brain barrier integrity during embryogenesis. Nature. 468, 562–6.

Diaz-Flores, L., et al., 2009. Pericytes. Morphofunction, interactions and pathology in a quiescent and activated mesenchymal cell niche. Histol Histopathol. 24, 909–69.

Diaz-Flores, L., et al., 1991. Microvascular pericytes: a review of their morphological and functional characteristics. Histol Histopathol. 6, 269–86.

Egginton, S., et al., 1996. In vivo pericyte-endothelial cell interaction during angiogenesis in adult cardiac and skeletal muscle. Microvasc Res. 51, 213–28.

Fang, J. S., et al., 2013. Connexin45 regulates endothelial-induced mesenchymal cell differentiation toward a mural cell phenotype. Arterioscler Thromb Vasc Biol. 33, 362–8.

Faraco, G., et al., 2017. Brain perivascular macrophages: characterization and functional roles in health and disease. J Mol Med (Berl). 95, 1143–1152.

Faraco, G., et al., 2016. Perivascular macrophages mediate the neurovascular and cognitive dysfunction associated with hypertension. J Clin Invest.

Fouquet, C., et al., 2015. Improving axial resolution in confocal microscopy with new high refractive index mounting media. PLoS One. 10, e0121096.

Gerhardt, H., et al., 2000. N-cadherin mediates pericytic-endothelial interaction during brain angiogenesis in the chicken. Dev Dyn. 218, 472–9.

Gonzales, A. L., et al., 2020. Contractile pericytes determine the direction of blood flow at capillary junctions. Proc Natl Acad Sci U S A. 117, 27022–27033.

Grant, R. I., et al., 2019. Organizational hierarchy and structural diversity of microvascular pericytes in adult mouse cortex. J Cereb Blood Flow Metab. 39, 411–425.

Hall, C. N., et al., 2014. Capillary pericytes regulate cerebral blood flow in health and disease. Nature. 508, 55–60.

Hammer, S., et al., 2015. Multiple Retinal Axons Converge onto Relay Cells in the Adult Mouse Thalamus. Cell Rep. 12, 1575–83.

Hartmann, D. A., et al., 2021. Brain capillary pericytes exert a substantial but slow influence on blood flow. Nat Neurosci. 24, 633–645.

Hill, R. A., et al., 2015. Regional Blood Flow in the Normal and Ischemic Brain Is Controlled by Arteriolar Smooth Muscle Cell Contractility and Not by Capillary Pericytes. Neuron. 87, 95–110.

Ivanova, E., et al., 2021. Retina-specific targeting of pericytes reveals structural diversity and enables control of capillary blood flow. J Comp Neurol. 529, 1121–1134.

Johnstone, S., et al., 2009. Biological and biophysical properties of vascular connexin channels. Int Rev Cell Mol Biol. 278, 69–118.

Leak, L. V., Burke, J. F., 1968. Ultrastructural studies on the lymphatic anchoring filaments. J Cell Biol. 36, 129–49.

Lee, S., et al., 2005. Processing of VEGF-A by matrix metalloproteinases regulates bioavailability and vascular patterning in tumors. J Cell Biol. 169, 681–91.

Lopez-Luppo, M., et al., 2017. Blood Vessel Basement Membrane Alterations in Human Retinal Microaneurysms During Aging. Invest Ophthalmol Vis Sci. 58, 1116–1131.

Maarouf, N., et al., 2017. Structural analysis of endothelial projections from mesenteric arteries. Microcirculation. 24.

Mayr, D., et al., 2021. Characterization of the two inducible Cre recombinase-based mouse models NG2-CreER(TM) and PDGFRb-P2A-CreER(T2) for pericyte labeling in the retina. Curr Eye Res.

Navarro, R., et al., 2016. Immune Regulation by Pericytes: Modulating Innate and Adaptive Immunity. Front Immunol. 7, 480.

Ornelas, S., et al., 2021. Three-dimensional ultrastructure of the brain pericyte-endothelial interface. J Cereb Blood Flow Metab. 271678X211012836.

Park, S. Y., et al., 2014. Cardiac, skeletal, and smooth muscle mitochondrial respiration: are all mitochondria created equal? Am J Physiol Heart Circ Physiol. 307, H346–52.

Payne, L. B., et al., 2020. Pericytes in Vascular Development. Current Tissue Microenvironment Reports. 1, 143–154.

Payne, L. B., et al., 2022. Pericyte Progenitor Coupling to the Emerging Endothelium During Vasculogenesis via Connexin 43. Arterioscler Thromb Vasc Biol. ATVBAHA121317324.

Payne, L. B., et al., 2019. The pericyte microenvironment during vascular development. Microcirculation. 26, e12554.

Petzold, G. C., Murthy, V. N., 2011. Role of astrocytes in neurovascular coupling. Neuron. 71, 782–97.

Potente, M., Makinen, T., 2017. Vascular heterogeneity and specialization in development and disease. Nat Rev Mol Cell Biol. 18, 477–494.

Rustenhoven, J., et al., 2017. Brain Pericytes As Mediators of Neuroinflammation. Trends Pharmacol Sci. 38, 291–304.

Sava, P., et al., 2015. Human microvascular pericyte basement membrane remodeling regulates neutrophil recruitment. Microcirculation. 22, 54–67.

Stratman, A. N., et al., 2009. Pericyte recruitment during vasculogenic tube assembly stimulates endothelial basement membrane matrix formation. Blood. 114, 5091–101.

Thakar, R. G., et al., 2009. Cell-shape regulation of smooth muscle cell proliferation. Biophys J. 96, 3423–32.

Thomsen, M. S., et al., 2017. The vascular basement membrane in the healthy and pathological brain. J Cereb Blood Flow Metab. 37, 3300–3317.

Tien, T., et al., 2014. Downregulation of Connexin 43 promotes vascular cell loss and excess permeability associated with the development of vascular lesions in the diabetic retina. Mol Vis. 20, 732–41.

Tillet, E., et al., 2005. N-cadherin deficiency impairs pericyte recruitment, and not endothelial differentiation or sprouting, in embryonic stem cell-derived angiogenesis. Exp Cell Res. 310, 392–400.

Tsilibary, E. C., 2003. Microvascular basement membranes in diabetes mellitus. J Pathol. 200, 537–46.

Volz, K. S., et al., 2015. Pericytes are progenitors for coronary artery smooth muscle. Elife. 4.

Wimmer, R. A., et al., 2019. Human blood vessel organoids as a model of diabetic vasculopathy. Nature. 565, 505–510.

Zhao, Z., et al., 2015. Establishment and Dysfunction of the Blood-Brain Barrier. Cell. 163, 1064–78.

