## Supplemental Information for "Pericyte Heterogeneity Identified by 3D Ultrastructural Analysis of the Microvessel Wall"

### Movie Legends

**Supplemental Movie 1.** Animation of a 360$^o$ rotation of the 3D rendering for the thin-strand pericyte shown in Figure 1. Blue represents the endothelial cell pseudo-coloring from corresponding SBF-SEM images (reduced opacity), and green represents the pericyte pseudo-coloring.

**Supplemental Movie 2.** Animation of a 360$^o$ rotation of the 3D rendering for the mesh pericyte shown in Figure 1. Blue represents the endothelial cell pseudo-coloring from corresponding SBF-SEM images (reduced opacity), and green represents the pericyte pseudo-coloring.

**Supplemental Movie 3.** Animation of a 360$^o$ rotation of the 3D rendering for the ensheathing pericyte shown in Figure 2. Blue represents the endothelial cell pseudo-coloring from corresponding SBF-SEM images, green represents the pericyte pseudo-coloring, and orange represents the medial cell pseudo-coloring.

**Supplemental Movie 4.** Animation of a 180$^o$ rotation horizontally and vertically of the 3D rendering for the peg-and-socket junction shown in Figure 4. Blue wire-frame represents the endothelial cell pseudo-coloring from corresponding SBF-SEM images, and green represents the pericyte pseudo-coloring.

**Supplemental Movie 5.** Animation of a 360$^o$ rotation of the 3D rendering for the medial cell-endothelial cell junctions (white arrowheads) shown in Figure 5. Blue wire-frame represents the endothelial cell pseudo-coloring from corresponding SBF-SEM images, green represents the pericyte pseudo-coloring, and orange represents the medial cell pseudo-coloring.

**Supplemental Movie 6.** Animation of a 360$^o$ rotation of the 3D rendering for the medial cell-endothelial cell junctions (white arrowheads) shown in Figure 5, and zoomed in from Supplemental Movie 5. Blue wire-frame represents the endothelial cell pseudo-coloring from corresponding SBF-SEM images, green represents the pericyte pseudo-coloring, and orange represents the medial cell pseudo-coloring.
